# Detection of pigeon circoviruses in ticks of sheep and camels in Inner Mongolia, China

**DOI:** 10.1101/2021.06.09.447674

**Authors:** Yunyi Kong, Chao Yan, Gang Zhang, Yurong Cai, Biao He, Yong Li

## Abstract

Circovirus is one of the smallest known DNA viruses infecting animals. In this report, circovirus was detected in two tick species- Hyalomma asiaticum and Dermacentor nuttalli collected from sheep and camels in Inner Mongolia, China by high-throughput sequencing. The reverse semi-nested PCR assay revealed a complete genomic sequence of 2042 base pairs of cyclic single-stranded DNA (ssDNA). Evolutionary analysis further revealed that it belonged to circovirus with two major open read frames (ORFs) that putatively encode replicase and capsid proteins, respectively. Sequence and phylogenetic analysis suggested that the circovirus belonged to pigeon circovirus. The results demonstrated the presence of pigeon circovirus in ticks in Inner Mongolia, which provide the basis for future exploration of the prevalence of pigeon circovirus in Inner Mongolia and the pathogens carried by ticks in Inner Mongolia.

## Introduction

Taxonomically, circovirus is a small, nonenveloped, single-stranded DNA virus with a circular genome, which is categorized into the genus Circovirus of the Circoviridae family (Rosario *et al.* 2012). Circoviral genomes range from 1.8 to 2.1 kb in size and contain two open reading frames that encode the replication (Rep) and capsid proteins (Cap) (Breitbart *et al.* 2017). Vertebrates, including birds, poultry and mammals, are natural hosts of circovirus. Circovirus infection may cause diseases in animals (Delwart and Li 2012). For example, porcine circovirus type 2 (PCV2) has been associated with various disease syndromes in pigs, primarily postweaning multisystemic wasting syndrome (PMWS) (Allan and Ellis 2000). Other symptoms in sows include abortion, dermatitis, and nephropathy syndrome (de Castro *et al.* 2012; Zhai *et al.* 2014; Kekarainen and Segalés 2015). Canine circovirus infection has also been reported to cause hemorrhagic enteritis that was associated with a sudden onset of weakened appetite, vomiting, and bloody diarrhea in dogs(Hsu *et al.* 2016; Dowgier *et al.* 2017). Similarly, avian circoviruses are related to symptoms of immunosuppression and feather abnormalities in birds or poultry(Todd 2000; Shearer *et al.* 2008; Zhang *et al.* 2009; Stenzel and Koncicki 2017).

Although most circoviruses currently cannot readily be cultured in vitro, with the exception that porcine circovirus (PCV) is able to grow in PK-15 cells(Stevenson *et al.* 1999), an increasing number of novel circovirus-like genomes have been discovered from wild mammals and insects using high-throughput sequencing techniques(Ge *et al.* 2011; Marton *et al.* 2015). Among them, however, there are few reports of circovirus in ticks, the second most common vectors of pathogens, despite porcine circovirus and raven-related circovirus being discovered in ticks(Wang *et al.* 2018; Franzo *et al.* 2019). To explore whether the ticks in Inner Mongolia autonomous region were the vectors of circovirus, we tried to detect circular ssDNA genomic sequences from ticks collected from sheep and camels in this location by high-throughput sequencing technology and the reverse semi-nested PCR.

## Materials and methods

A total of 2,662 ticks were collected from sheep and camels in the Alxa Left Banner, Alxa Right Banner and Siziwang Banners of Inner Mongolia, China, from April to May 2016. For each sampling site, three flocks of sheep or camels were selected, and more than 50 animals in each flock were randomly selected for sampling. Ticks on the skin surface of animals were caught and collected in a container. All samples were frozen and stored at −80 °C. The collected ticks were pooled according to the hosts they were collected from and the species of ticks. Approximately 15 ticks were pooled in one sample for processing and analysis. To prepare samples, ticks of each pool were put into a 2-mL centrifuge tube containing 1 mL of Dulbecco’s modified Eagle’s medium (DMEM; Gibco, Carlsbad, USA) prior to being homogenized in a frozen grinder (Jingxin, Shanghai, China). The homogenized samples were then centrifuged at 4 °C and 12,000× g for 10 min. The supernatants were collected, and total RNA was extracted from tick pools with an RNeasy Mini Kit (QIAGEN, Hiden, Germany), followed by cDNA synthesis using M-MLV reverse transcriptase (TaKaRa Biotechnology, Dalian, China). The qualified cDNAs were subjected to high-throughput sequencing for viral metagenomic analysis by the Beijing Genome Institute (BGI, Shenzhen, China).

After removing host-derived sequences, 3,568,561 reads with an average length of 145 nt were obtained, of which 12,931 (0.004%) were annotated to viruses of 20 families and some unclassified viruses, including dsDNA, ssDNA, and ssRNA viruses (Supplemental Fig. 1). According to the sequencing results, the DNA of each tick pool was extracted using a TIANamp Genomic DNA Kit (Tiangen, Beijing, China), and the fractional genes of circovirus were determined by PCR using the primers TiCV-F1, TiCV-F2 and TiCV-R (Supplemental Table 1). The viral metagenomic results were also corroborated by a PCR assay. To obtain the complete sequence of circovirus, primers for amplifying the complete genome were designed according to the results of high-throughput sequencing (Supplemental Table 2). The PCR was performed in a 50 μL reaction system containing 1 μL of genomic DNA as a template from each pool sample, 25 μL of 2× Phabta Max Master Mix (Vazyme, Nanjing, China), 20 μL of ddH2O, and 2 μL each of the forward and reverse primers (10 μM). The PCRs were conducted in a thermal cycler (Analytik Jena, Thuringia, Germany) using the following thermocycling conditions: predenaturation at 95◻°C for 3◻min, followed by 35 cycles (denaturation for 15◻s at 95◻°C, annealing for 15◻s at 57◻°C, and extension for 15-47◻s at 72◻°C), and a final incubation for 5◻min at 72◻°C. The PCR products were analyzed by electrophoresis using a 1.0% agarose gel stained with ethidium bromide.

All amplified products were analyzed by Sanger sequencing (ComateBio, Changchun, China). Nucleotide sequence homology was performed by BLAST analysis (http://www.ncbi.nlmn.nih.gov/BLAST). All the sequencing results were assembled and aligned using CLUSTAL W in MEGA 7.0 (http://www.megasoftware.net/) and compared with the sequences of other circovirus strains in GenBank. A phylogenetic tree was constructed with the maximum-likelihood method by M EGA 7.0 software. The reliability of the branches of the tree was assessed using a bootstrap analysis with 1,000 replicates.

## Results

All samples were screened for the presence of circovirus by half-nested PCR. The distribution of circovirus in these three regions was assessed, and circoviral DNA was detected in tick samples from Alxa Left Banner and Siziwang Banner but not in samples from Alxa Right Banner. The positive percentages of circoviral DNA were 0.093% in ticks from sheep and 1.268% in ticks from camels in Alxa Left Banner, Inner Mongolia and 3.734% in ticks from sheep in Siziwang Banner, Inner Mongolia. The positive rate was determined by calculating the minimum infection rate (MIR), which was the ratio of the number of positive tick pools to the total number of analyzed ticks(Gu *et al.* 2003). The analysis data are listed in Table 1.

**Table 1.**
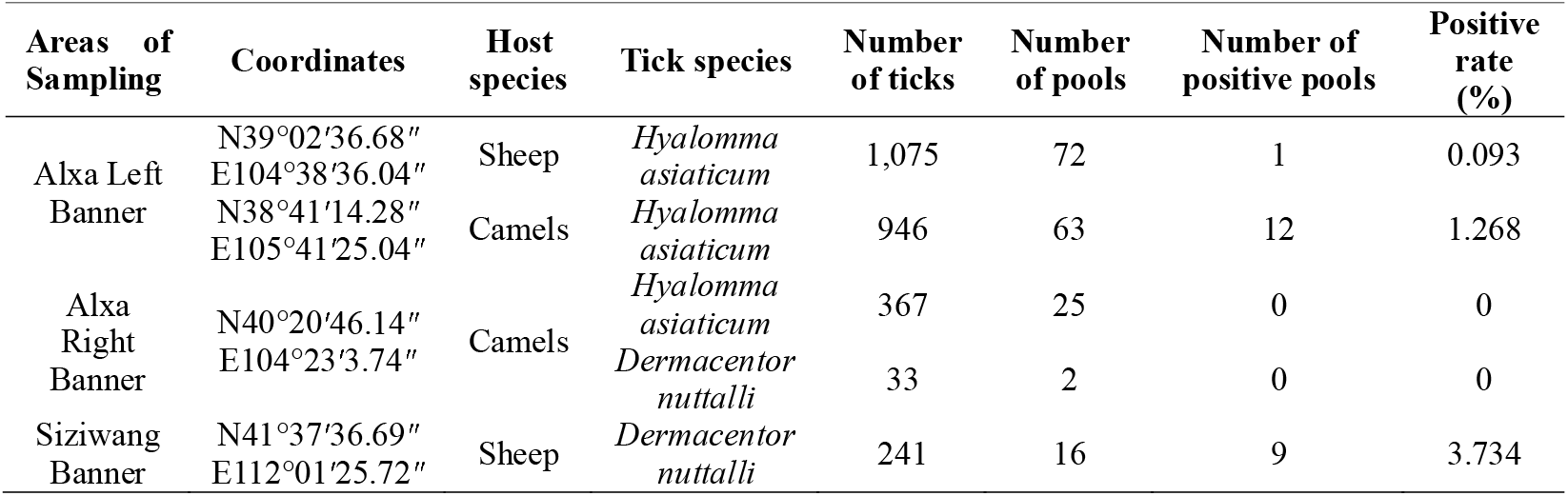
Molecular detection of circovirus from ticks isolated from different regions in Inner Mongolia, China

The complete sequences of circoviruses recovered by half-nested PCR using the specific primer sets based on Sanger sequencing revealed a circular viral genome of 2,042 nucleotides (nt) in ticks. The genomic sequence obtained in this study was submitted to GenBank with accession number MN920392.

To explore the evolutionary relationship between this circovirus and other circoviruses, the complete genomic sequences and ORFs of circoviruses were used for phylogenetic tree mapping and homology analysis. In this report, genome sequences of 19 members of this family were analyzed. The phylogenetic analysis of the complete viral genome showed that this virus was close to pigeon circovirus (PiCV) (Fig. 1 A). In addition, the phylogeny also suggested that tick circovirus (TiCV) has a common ancestor with the evolutionary branches of starling circovirus (StCV), raven circovirus (RaCV), finch circovirus (FiCV) and zebra finch circovirus. Except for chimpanzee feces-associated circovirus (accession no. GQ404851), mammalian and avian circoviruses were relatively independent of the evolutionary tree. The phylogenetic trees of Rep and Cap were consistent with the whole genome (Fig 2).

**Fig. 1.**
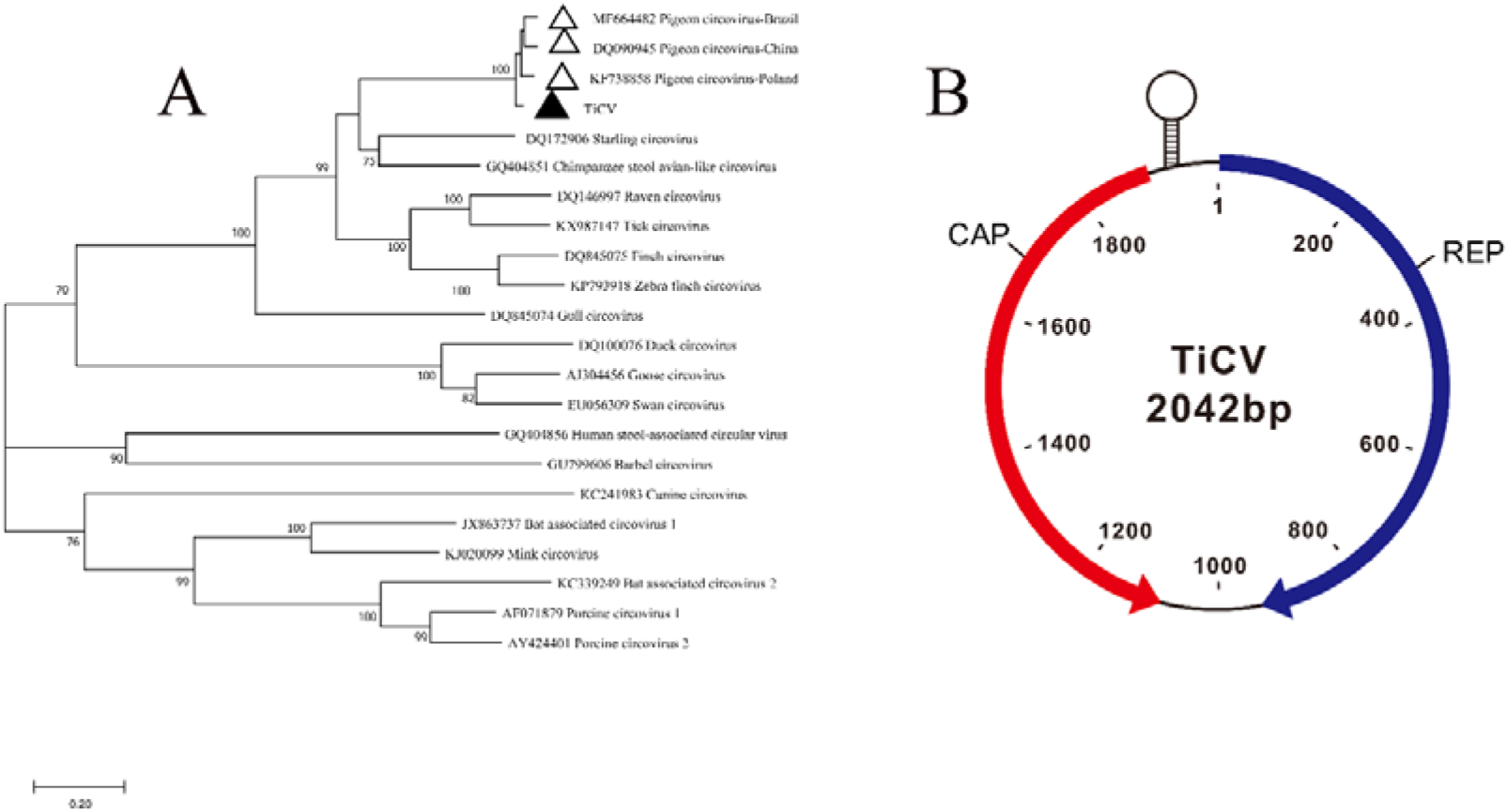
Phylogenetic tree and schematic representation of circovirus in ticks. A. Phylogenetic tree of the tick circovirus (TiCV) complete sequence. The tree was constructed using the maximum-likelihood method with MEGA 7.0 software. Bootstrap testing (1,000 replicates) was performed, and values lower than 60% are not shown. The bar at the bottom of the figure denotes the distance. B. Genome map of TiCV showing the ORFs encoding Rep and Cap.

**Fig. 2.**
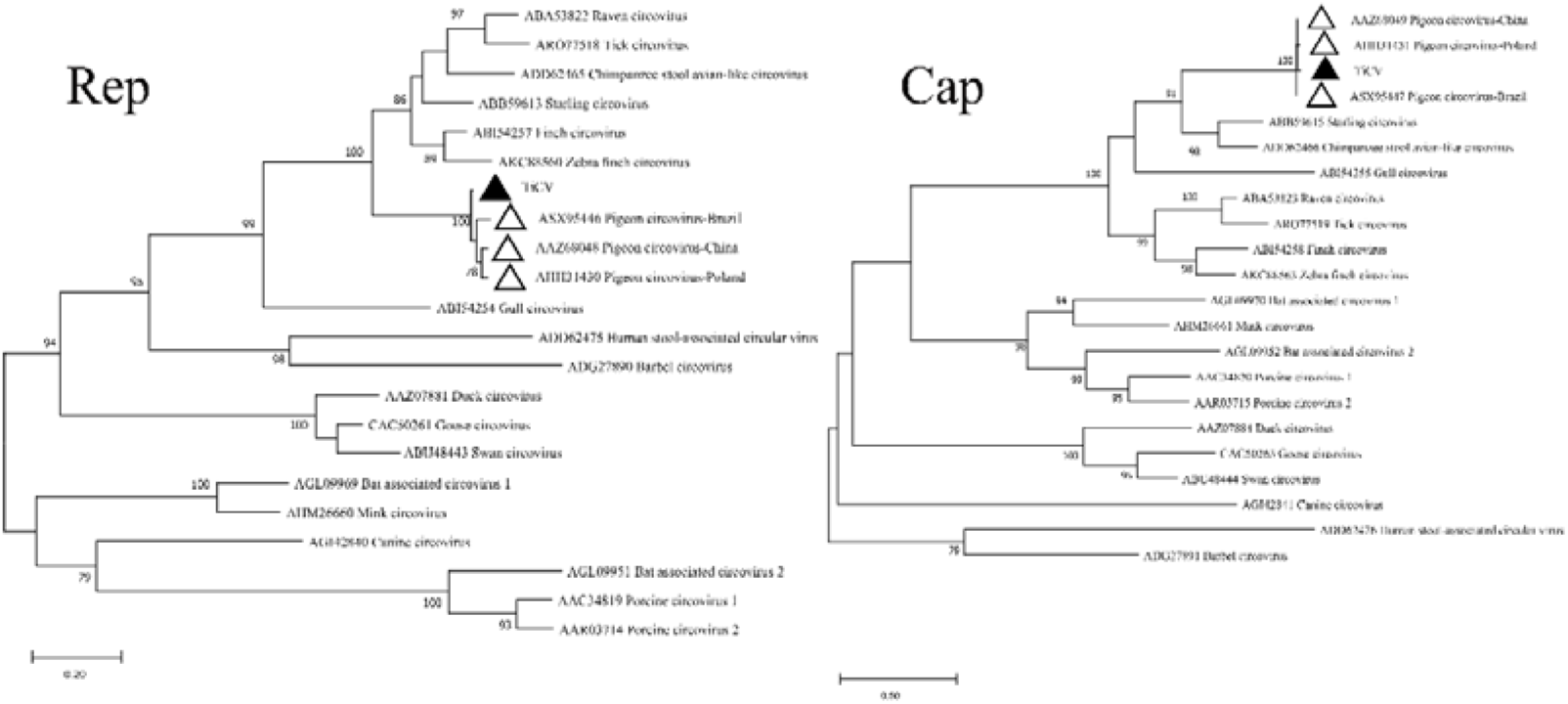
Phylogenetic tree of Tick circovirus (TiCV) based on Rep and Cap sequences

Through comparison of the TiCV genome with those of other circoviruses, a high identity of 93.5% to 95.5% with the genome of PiCV and a low identity (42.7.0% to 59.8%) with duck circovirus (DuCV), goose circovirus (GoCV), gull circovirus (GuCV), RaCV, FiCV and StCV were found (Supplemental Table 2). There were two large ORFs for this circovirus, which encode the replication-associated proteins (Rep) and the viral capsid protein (Cap). The intergenic region between the two ORFs could form a potential stem-loop structure, according to the mfold web server (Fig. 1 B). The putative Rep protein of TiCV was 315 amino acids (aa), and it had a high identity with the Rep proteins of FiCV (75.2%), RaCV (74.5%), StCV (73.3%), and PiCVs that were isolated from China (96.1%) and Brazil (97.1%). It had a low identity with those of GuCV (61.3%), human stool-associated circovirus (55.9%), DuCV (50.9%), and GoCV (51.9%) (Table 2). The putative Cap protein of TiCV was 273 aa and shared a higher identity with that of PiCVs from China (98.9%) and Brazil (98.6%) than that of with other circoviruses, including FiCV (44.9%), Tick cirocvirus (41.0%), and StCV (60.1%) (Table 2).

**Table 2.** Comparison of amino acid and nucleotide sequence identity between tick circovirus and other circoviruses

| Viral name | GenBank<br>accession number | Rep segment of<br>TiCV |  | Cap segment of<br>TiCV |  |
| --- | --- | --- | --- | --- | --- |
|  |  | nt (%) | aa (%) | nt (%) | aa (%) |
| Porcine circovirus 1 | AF071879 | 49.4 | 43.5 | 37.7 | 39.7 |
| Goose circovirus | AJ304456 | 54.2 | 51.9 | 40.5 | 35.4 |
| Porcine circovirus 2 | AY424401 | 48.8 | 43.8 | 37.7 | 39.3 |
| Tick circovirus | KX987147 | 56.6 | 71.8 | 49.4 | 41.0 |
| Duck circovirus | DQ100076 | 56.2 | 50.7 | 36.4 | 31.9 |
| Raven circovirus | DQ146997 | 72.8 | 74.5 | 54.6 | 48.3 |
| Starling circovirus | DQ172906 | 72.2 | 73.3 | 62.5 | 60.1 |
| Gull circovirus | DQ845074 | 64.9 | 61.3 | 52.2 | 56.4 |
| Finch circovirus | DQ845075 | 74.2 | 75.2 | 54.5 | 44.9 |
| Swan circovirus | EU056309 | 53.4 | 51.2 | 40.3 | 37.7 |
| Chimpanzee feces-associated<br>circovirus | GQ404851 | 72.4 | 74.8 | 63.2 | 66.5 |
| Human stool-associated circular<br>virus | GQ404856 | 53.1 | 55.9 | 32.9 | 30.4 |
| Barbel circovirus | GU799606 | 51.6 | 51.2 | 37.3 | 34.2 |
| Bat associated circovirus 1 | JX863737 | 51.3 | 47.8 | 36.6 | 36.6 |
| Canine circovirus | KC241983 | 53.7 | 45.7 | 29.4 | 23.3 |
| Bat associated circovirus 2 | KC339249 | 48.6 | 44.1 | 38.4 | 42.4 |
| Mink circovirus | KJ020099 | 52.9 | 47.8 | 34.5 | 38.1 |
| Zebra finch circovirus | KP793918 | 71.6 | 75.3 | 52.5 | 54.9 |
| Pigeon circovirus-zj1 | DQ090945 | 94.9 | 96.0 | 95.4 | 98.9 |
| Pigeon circovirus-PL44B | KF738858 | 94.7 | 96.6 | 95.6 | 98.0 |
| Pigeon circovirus- SLMA4 | MF664482 | 94.6 | 97.1 | 95.8 | 98.6 |

## Discussion

Ticks are efficient vectors for the transmission of pathogens, including many arboviruses. To our knowledge, the identified tick-borne viruses include members of two orders, nine families and at least twelve genera(Shi *et al.* 2018). The infection of most of these tick-borne viruses may cause diseases in humans or animals(Vandegrift and Kapoor 2019). With a vast territory and well-balanced ecological environment in Inner Mongolia, it relies heavily on the animal husbandry industry. In this report, we attempted to detect circular ssDNA genomic sequences from ticks collected from sheep and camels in Inner Mongolia, China, by high-throughput sequencing technology.

Notably, genomic DNA of PiCV was detected in ticks parasitized on sheep and camels at two remote sampling sites in Inner Mongolia, suggesting that PiCV exists in ticks in Inner Mongolia. Interestingly, however, the ticks were collected from sheep and camels herding on the desert steppe or meadow grassland in Inner Mongolia, where no pigeons live.

In addition, circoviruses may be able to transmit between different species. It has been reported that a PiCV was detected in chicken and that a PCV was detected in cow or calf samples(Li *et al.* 2010; Li *et al.* 2011). However, there is no direct evidence that this virus can infect ruminants, such as sheep or camels. These findings suggest that we can establish a serological detection method using antibodies against circovirus in further research, combined with nucleic acid detection methods, which may solve this problem. Notably, the role played by ticks in the transmission route of circovirus is currently uncertain. Whether ticks are used as a source of transmission to spread the virus to the host or as an intermediate host from an infected animal to a tick is worthy of additional experiments, such as a serological detection of circovirus, for further validation(Estrada-Pena and de la Fuente 2014). Nevertheless, the existence of genomic DNA of PiCV was detected in ticks of sheep and camels in Inner Mongolia, China.

## Conclusions

In this study, we detected for the first time that the PiCV was found in ticks of sheep and camels in Inner Mongolia, China. Sequence comparison and phylogenetic analysis further identified that the new circovirus sequence belongs to PiCV and was detected in *Hyalomma asiaticum* and *Dermacentor nuttalli*. This study thus made a preliminary report on the genome structure and epidemic situation of circovirus in ticks of sheep and camels in Inner Mongolia, China, for the first time.

## Author contributions

YL designed the research. YYK, CY, GZ, YRC and BH helped with the experiments, data analysis, and discussion. YYK and YL wrote the manuscript.

## Compliance with ethical standards

### Ethics approval and consent to participate

The experiments involving sheep and camels were performed according to protocols approved by the Institutional Animal Care and Use Committee of Ningxia University (NXU-2019-007). The need for consent is deemed unnecessary according to national regulations, but an informed verbal consent was obtained from the sheep and camel owners. The ethics committee for the use of animals of Ningxia University approved this study.

### Informed consent

Informed consent was obtained from all individual participants in the study.

---

**Supplemental Table 1.**
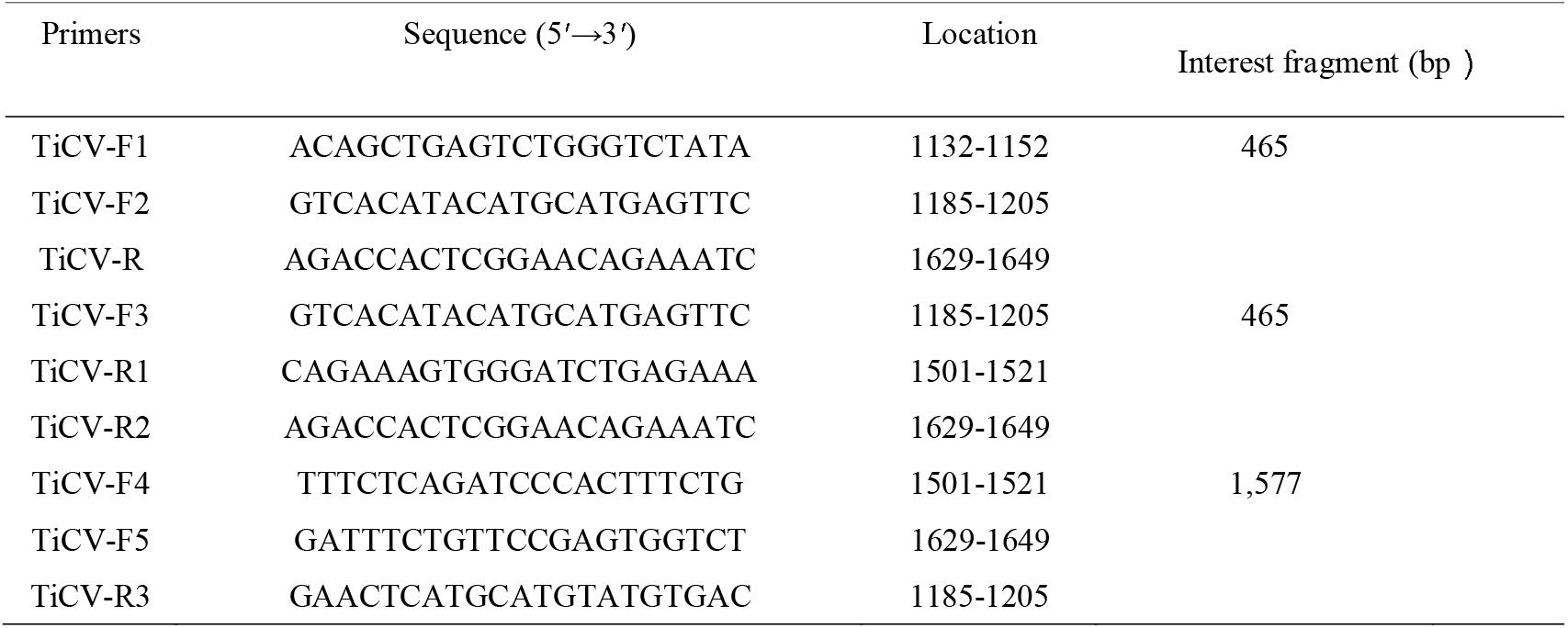
Oligonucleotide primers for amplifying the complete genome of circovirus in ticks

**Supplemental Table 2.** Full-length sequence comparison of tick circoviruses with others

| Viral name | GenBank accession number | TiCV |
| --- | --- | --- |
|  |  | nt (%) |
| Porcine circovirus 1 | AF071879 | 42.7 |
| Goose circovirus | AJ304456 | 43.3 |
| Porcine circovirus 2 | AY424401 | 42.5 |
| Duck circovirus | DQ100076 | 42.7 |
| Raven circovirus | DQ146997 | 57.4 |
| Finch circovirus | DQ845075 | 57.3 |
| Starling circovirus | DQ172906 | 59.8 |
| Gull circovirus | DQ845074 | 52.1 |
| Swan circovirus | EU056309 | 42.7 |
| Chimpanzee feces-associated circovirus | GQ404851 | 65.3 |
| Human stool-associated circular virus | GQ404856 | 39.9 |
| Barbel circovirus | GU799606 | 42.1 |
| Bat associated circovirus 1 | JX863737 | 42.5 |
| Canine circovirus | KC241983 | 43.2 |
| Bat associated circovirus 2 | KC339249 | 42.6 |
| Mink circovirus | KJ020099 | 42.2 |
| Zebra finch circovirus | KP793918 | 57.1 |
| Tick circovirus | KX987147 | 56.5 |
| Pigeon circovirus-PL44B | KF738858 | 94.7 |
| Pigeon circovirus-zj1 | DQ090945 | 93.5 |
| Pigeon circovirus- SLMA4 | MF664482 | 95.5 |

**Supplementary Fig. 1.**
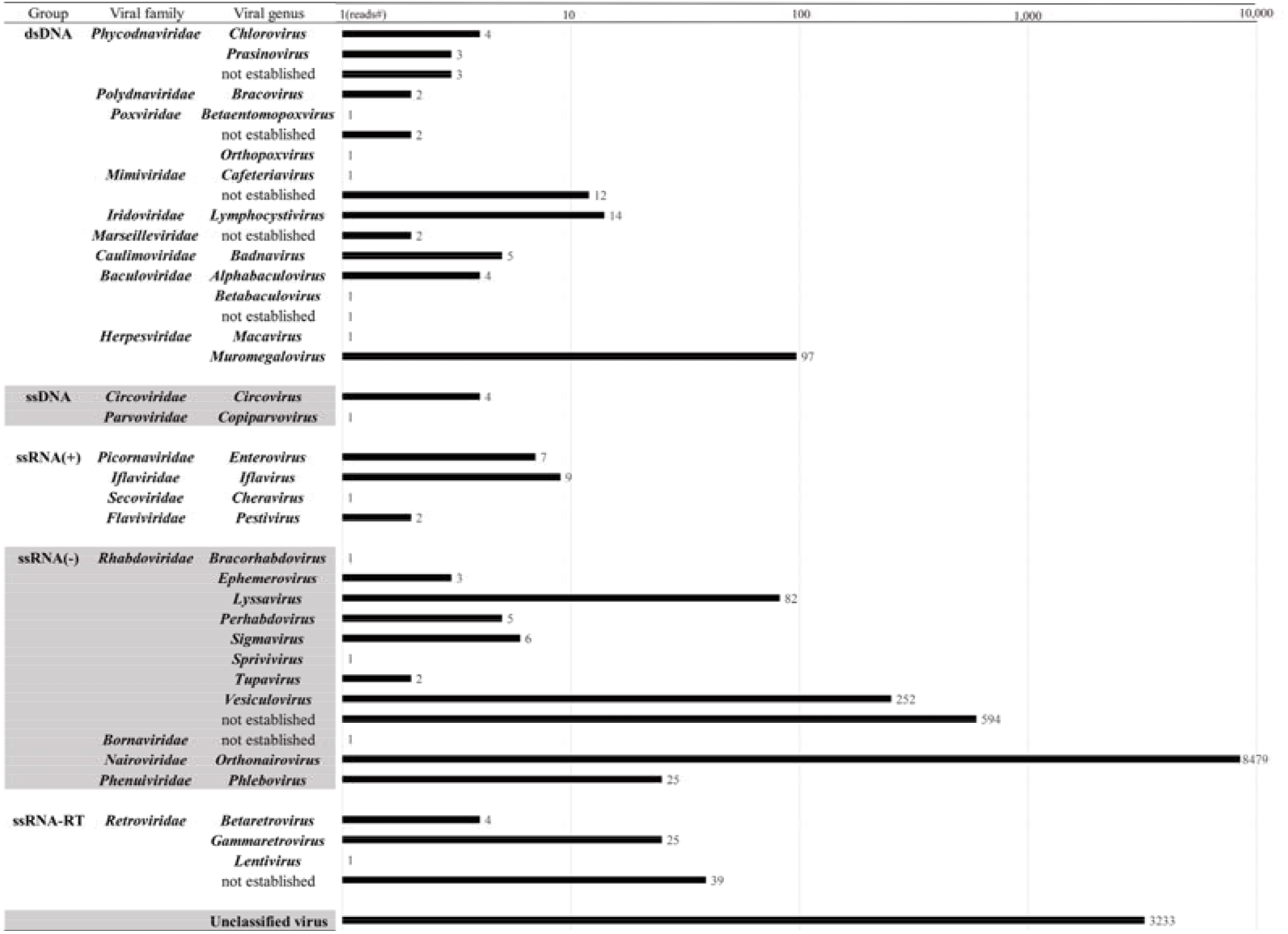
Numbers of reads annotated to viruses classified by viral genus, family and nucleic acid group.

## References

Allan GM, Ellis JA (2000) Porcine circoviruses: a review. J Vet Diagn Invest 12:3–14

Breitbart M, Delwart E, Rosario K, Segales J, Varsani A, Ictv Report C (2017) ICTV Virus Taxonomy Profile: Circoviridae. J Gen Virol 98:1997–1998

de Castro AM, Cruz TF, Salgado VR, Kanashiro TM, Ferrari KL, Araujo JP, Brandão PE, Richtzenhain LJ (2012) Detection of porcine circovirus genotypes 2a and 2b in aborted foetuses from infected swine herds in the State of São Paulo, Brazil. Acta Veterinaria Scandinavica 54:29

Delwart E, Li L (2012) Rapidly expanding genetic diversity and host range of the Circoviridae viral family and other Rep encoding small circular ssDNA genomes. Virus Res 164:114–121

Dowgier G, Lorusso E, Decaro N, Desario C, Mari V, Lucente MS, Lanave G, Buonavoglia C, Elia G (2017) A molecular survey for selected viral enteropathogens revealed a limited role of Canine circovirus in the development of canine acute gastroenteritis. Vet Microbiol 204:54–58

Estrada-Pena A, de la Fuente J (2014) The ecology of ticks and epidemiology of tick-borne viral diseases. Antiviral Res 108:104–128

Franzo G, Grassi L, Tucciarone CM, Drigo M, Martini M, Pasotto D, Mondin A, Menandro ML (2019) A wild circulation: High presence of Porcine circovirus 3 in different mammalian wild hosts and ticks. Transbound Emerg Dis 66:1548–1557

Ge X, Li J, Peng C, Wu L, Yang X, Wu Y, Zhang Y, Shi Z (2011) Genetic diversity of novel circular ssDNA viruses in bats in China. J Gen Virol 92:2646–2653

Gu WD, Lampman R, Novak RJ (2003) Problems in estimating mosquito infection rates using minimum infection rate. Journal of Medical Entomology 40:595–596

Hsu HS, Lin TH, Wu HY, Lin LS, Chung CS, Chiou MT, Lin CN (2016) High detection rate of dog circovirus in diarrheal dogs. BMC Vet Res 12:116

Kekarainen T, Segalés J (2015) Porcine circovirus 2 immunology and viral evolution. Porcine Health Management 1

Li L, Kapoor A, Slikas B, Bamidele OS, Wang C, Shaukat S, Masroor MA, Wilson ML, Ndjango JB, Peeters M, Gross-Camp ND, Muller MN, Hahn BH, Wolfe ND, Triki H, Bartkus J, Zaidi SZ, Delwart E (2010) Multiple diverse circoviruses infect farm animals and are commonly found in human and chimpanzee feces. J Virol 84:1674–1682

Li L, Shan T, Soji OB, Alam MM, Kunz TH, Zaidi SZ, Delwart E (2011) Possible cross-species transmission of circoviruses and cycloviruses among farm animals. J Gen Virol 92:768–772

Marton S, Ihasz K, Lengyel G, Farkas SL, Dan A, Paulus P, Banyai K, Feher E (2015) Ubiquiter circovirus sequences raise challenges in laboratory diagnosis: the case of honey bee and bee mite, reptiles, and free living amoebae. Acta Microbiol Immunol Hung 62:57–73

Rosario K, Duffy S, Breitbart M (2012) A field guide to eukaryotic circular single-stranded DNA viruses: insights gained from metagenomics. Arch Virol 157:1851–1871

Shearer PL, Bonne N, Clark P, Sharp M, Raidal SR (2008) Beak and feather disease virus infection in cockatiels (Nymphicus hollandicus). Avian Pathol 37:75–81

Shi J, Hu Z, Deng F, Shen S (2018) Tick-Borne Viruses. Virol Sin 33:21–43

Stenzel T, Koncicki A (2017) The epidemiology, molecular characterization and clinical pathology of circovirus infections in pigeons – current knowledge. Vet Q 37:166–174

Stevenson GW, Kiupel M, Mittal SK, Kanitz CL (1999) Ultrastructure of porcine circovirus in persistently infected PK-15 cells. Vet Pathol 36:368–378

Todd D (2000) Circoviruses: immunosuppressive threats to avian species: a review. Avian Pathol 29:373–394

Vandegrift KJ, Kapoor A (2019) The Ecology of New Constituents of the Tick Virome and Their Relevance to Public Health. Viruses 11

Wang B, Sun LD, Liu HH, Wang ZD, Zhao YK, Wang W, Liu Q (2018) Molecular detection of novel circoviruses in ticks in northeastern China. Ticks Tick Borne Dis 9:836–839

Zhai SL, Chen SN, Xu ZH, Tang MH, Wang FG, Li XJ, Sun BB, Deng SF, Hu J, Lv DH, Wen XH, Yuan J, Luo ML, Wei WK (2014) Porcine circovirus type 2 in China: an update on and insights to its prevalence and control. Virol J 11:88

Zhang X, Jiang S, Wu J, Zhao Q, Sun Y, Kong Y, Li X, Yao M, Chai T (2009) An investigation of duck circovirus and co-infection in Cherry Valley ducks in Shandong Province, China. Vet Microbiol 133:252–256

